# Mitochondrial dynamics regulate cell size in the developing cochlea

**DOI:** 10.1101/2024.03.04.583298

**Authors:** James D. B. O’Sullivan, Stephen Terry, Claire A. Scott, Anwen Bullen, Daniel J. Jagger, Zoë F. Mann

## Abstract

In multicellular tissues, cell size and shape are intricately linked with physiological function. In the vertebrate auditory organ, the neurosensory epithelium develops as a mosaic of sensory hair cells (HCs), and their glial-like supporting cells (SCs), which have distinct morphologies at different frequency positions along its tonotopic long axis. In the chick cochlea, the basilar papilla (BP), proximal (high-frequency) HCs are larger than their distal (low-frequency) counterparts, a morphological feature essential for frequency tuning. Mitochondrial dynamics, which constitute the equilibrium between fusion and fission, regulate differentiation and functional refinement across a variety of cell types. We investigate this a potential mechanism for cell size regulation in developing HCs. Using live imaging in intact BP explants, we identify distinct remodelling of mitochondrial networks in proximal compared to distal HCs. Manipulating mitochondrial dynamics in developing HCs alters their normal morphology along the proximal-distal (tonotopic) axis. Inhibition of the mitochondrial fusion machinery decreased proximal HC size, whilst promotion of fusion increased the distal HC size. We identify mitochondrial dynamics as a key regulator of HC size and morphology in developing inner ear epithelia.

**Summary Statement:** Mitochondrial remodelling drives developmental changes in cell size in the auditory sensory epithelium. Our data reveal a fundamental mechanism regulating cell size and frequency-place coding in the developing cochlea.

## Introduction

A unique feature of the avian cochlear epithelium is that the position of a hair cell (HC) along its proximal-distal long axis determines the sound frequency to which it will respond, a phenomenon known as “tonotopy” (Mann and Kelley, 2011).The tonotopic patterning of the avian cochlea (BP) is characterised in part by a proximal-to-distal gradient in HC size (Tilney and Saunders, 1983) (Fig 1A). High-frequency HCs at the proximal end of the BP are significantly larger that the low-frequency distal HCs at the opposite end. Specification of a HC’s ‘tonotopic identity’ relies on the coordinated signalling between graded glucose metabolism and morphogen signalling prior to their terminal differentiation at around embryonic day (E) 6-8 (Mann et al., 2014; Thiede et al., 2014; Son et al., 2015; O’Sullivan et al., 2023). Once instructed of their tonotopic identity, little is known about the effector mechanisms that generate the size differences between proximal and distal HCs in the mature epithelium.

**Figure 1:**
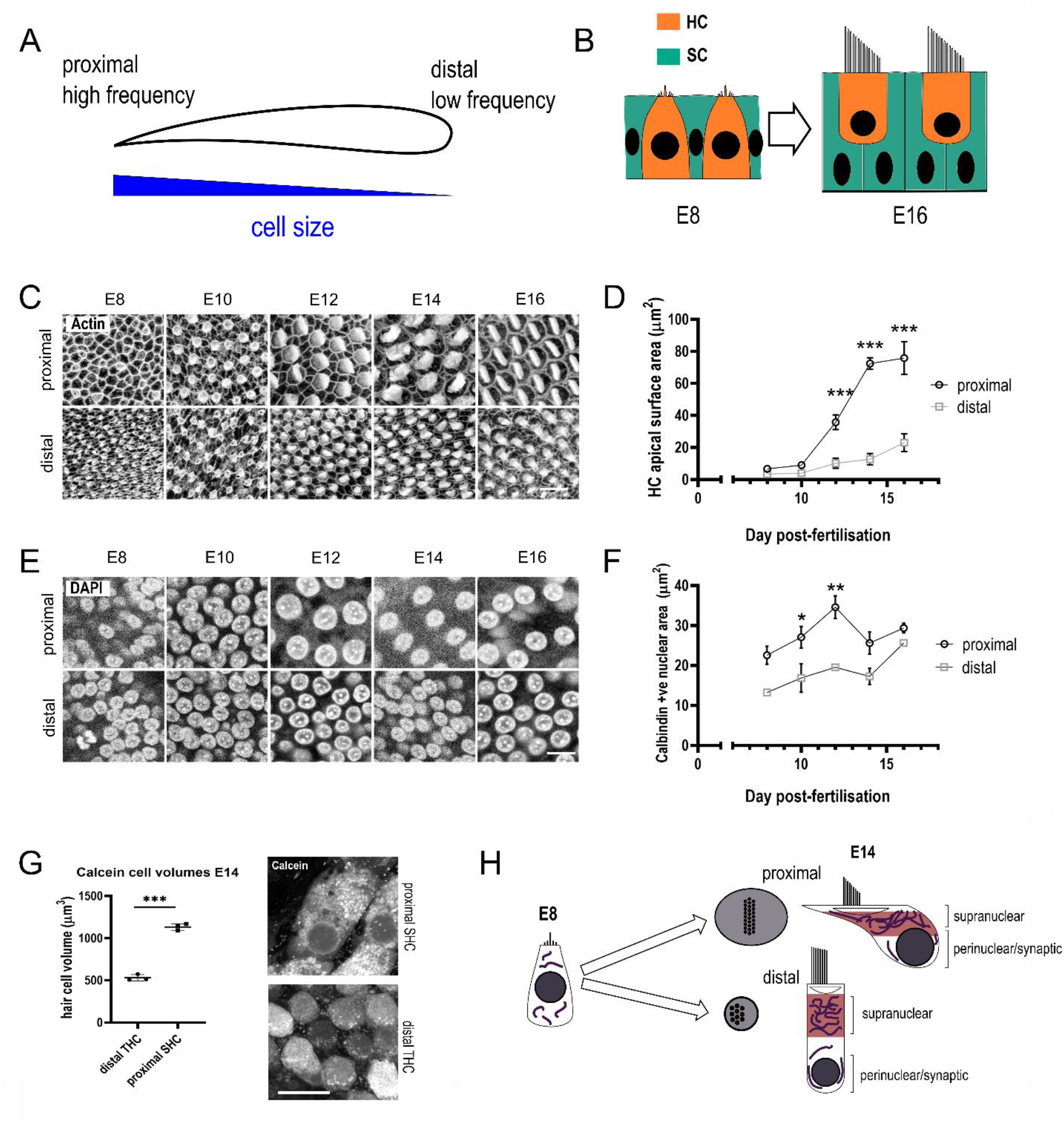
Hair cell size and shape changes in the developing cochlea. **(A)** Illustration of the avian auditory sensory epithelium. Hair Cells (HCs) are arranged longitudinally along the organ depending on their frequency sensitivity, and this arrangement (tonotopy) is correlated with a gradient in HC size. **(B)** Changes in the structure of the auditory sensory epithelium between E8 and E14. HCs are surrounded by a mosaic of non-sensory supporting cells (SCs). **(C)** Whole mount phalloidin staining of the proximal and distal BP HCs between E8-E16. Scale = 20 µm. **(D)** Quantification of HC luminal surface area in proximal and distal regions between E8 – E16. Data are Mean ± SD, n = 3 embryos, 15-40 HCs per embryo, ^***^ p ≤ 0.001, 2-way ANOVA, Sidak’s multiple comparisons, F = 1354, DF = 3. **(E)** Whole mount DAPI staining of proximal and distal HCs between E8-E16. Scale = 10 μm. **(F)** Quantification of HC nuclear area between E8 – E16. Data are Mean ± SD, n = 3 embryos, n = 15-40 HCs per embryo, ^*^ p > 0.05, ^**^ p > 0.01. 2-way ANOVA, Sidak’s multiple comparisons, F = 45.74, DF = 3. **(G)** Live imaging of calcein-AM loaded E14 explants from which HC volume was estimated using stereology. Data are mean ± SD, n=3, ^***^ p < 0.001. Unpaired t-test, t = 19.03, DF = 4. **(H)** Illustration of changes in HC morphology between E8 and E14 in proximal and distal BP regions. The supranuclear region of HCs (red), which in proximal HCs undergo a burst of growth between E10 and E14 in their supranuclear relative to the perinuclear region. The supranuclear region contains a high density of mitochondria.

Mitochondrial dynamics, referring to the distribution of mitochondria throughout a cell and remodelling of their networks by fusion and fission, have emerged as a key determinant of cell size and function (Baum and Gama, 2021; Fu et al., 2019). The dynamic equilibrium between fusion and fission also determines the morphological and metabolic properties of individual mitochondria. Biases of the mitochondrial population towards fission or fusion are unique to different cell types and tailored to support their specific functions. Fusion and fission are coordinated by a set of mitochondrial proteins (Van Der Bliek et al., 2013). The Mitofusins 1 and 2 (MFN1 and MFN2) regulate dynamics of the outer mitochondrial membranes (OMMs) (Chen et al., 2003), Optic Atrophy 1 (OPA1) coordinates crista structure and inner membrane (IMM) fusion (Patten et al., 2014; Wong et al., 2000), and Dynamin 1-like (DNM1L; Drp1 in mammals) separates individual mitochondria through the formation of a contractile ring (Chang et al., 2010; Ji et al., 2017). Asymmetric fission regulates mitochondrial quality control by segregating compromised mitochondria from the rest of the network and targeting them for removal via mitophagy (Song et al., 2015). Fusion promotes mitochondrial longevity and lessens the energetic load imposed on a single organelle (Fu et al., 2019).

Mounting evidence implicates mitochondrial dynamics as a regulator of the signalling pathways driving cell fate decisions during development (Khacho et al., 2016). For example, fission regulates differentiation in neural progenitors and morphological remodelling of T-cells during activation (Cervantes-Silva et al., 2021; Iwata et al., 2020). Increased mitochondrial fusion is important for specifying the distinct morphologies of neurons and astrocytes during neural development (Zehnder et al., 2021). An increase in mitochondrial biogenesis and fusion has long been considered a consequence of the higher metabolic demand posed by larger cells (Kitami et al., 2012; Posakony et al., 1977), a hypothesis supported by work showing that biasing cells towards fusion increases their overall size (Miettinen and Björklund, 2016).

Here, we use the tonotopic gradient in cell size (Fig 1A) along the chick BP as a model in which to interrogate the metabolic mechanisms regulating cell size. We find that mitochondrial dynamics and network architecture are extensively remodelled in BP HCs throughout development, and that biasing towards fusion or fission within a critical window of development alters HC size and tonotopic identity.

## Results & Discussion

Changes in HC size in proximal and distal regions of the BP were characterised between E8, when the majority of HCs are terminally differentiated, and at E16, when HCs are morphologically and functionally mature (Fig 1B) (Lippe, 1995). We observed no difference in HC lumenal surface area (LSA), an indicator of HC size, at E8 or E10, but differential growth along the tonotopic axis between E10 and E14 (Fig 1C, D). These size differences plateaued between E14 and E16, identifying E14 as a potential growth endpoint. In contrast to LSA, proximal calbindin-positive HCs had larger nuclei compared to distal HCs at E10, although this size difference was lost by E14 (Fig 1E, F). Contrary to previous studies in HCs from the mammalian cochlea (Fettiplace and Nam, 2019), 3D imaging of HCs from live BP explants revealed that by E14 (growth endpoint), the volume of proximal HCs is roughly twice that of distal HCs (Fig 1G).Therefore, the high ratio of LSA growth compared to nuclear growth and overall cell volume indicates that the area above the HC nucleus (hereafter referred to as supranuclear region) exhibits accelerated growth compared to the rest of the cell during development (Fig 1 H).

HCs contain a variety of spatially distinct mitochondrial sub-populations (Fig S1A), the majority residing in the supranuclear region towards the apical portion of the cell. Given the asymmetric growth of HCs, we sought to investigate how the properties of mitochondria in the supranuclear region change from E8 to E14. Transmission electron microscopy (TEM) in proximal and distal HCs revealed that mitochondria became thicker and acquired denser laminar cristae arrangements between E8 and E14 (Fig S1B), indicating a higher capacity for oxidative phosphorylation (Cogliati et al., 2013). To determine whether altered mitochondrial dynamics correlate with the proximal-distal gradient in supranuclear growth, we investigated remodelling of mitochondrial networks in developing proximal and distal HCs.

To visualise changes in mitochondrial morphology throughout development, live cochlear explants were loaded with the fluorescent mitochondrial dye tetramethyl rhodamine methyl ester (TMRM) between E8 and E14 (Fig 2), encompassing all stages of HC maturation (Si et al., 2003). Raw fluorescence intensity data were processed and segmented to generate 3D mitochondrial networks (Fig S3). Principle Component Analysis (PCA) (Wiemerslage and Lee, 2016) revealed that HC mitochondria developed along distinct morphological trajectories in proximal and distal regions (Fig 2A - D, Fig S1C). Mitochondrial morphology in both regions was comparable at E8, indicated by the high overlap of PCA clusters (Fig S1C). Mitochondrial volume in proximal HCs increased between E8 and E14 (Fig 2D), suggesting greater mitochondrial fusion and overall network growth in the high frequency region (Miettinen and Björklund, 2016). At E10, proximal and distal mitochondria were segregated largely based on differences in branch thickness (Fig 2D, Fig S1C). Thicker mitochondrial branches accommodate more lamellar cristae, which increases their capacity for ATP production (Afzal et al., 2021). The thicker mitochondrial branches in distal HCs at E10 may therefore reflect a more differentiated metabolic state, given the distal-to-proximal gradient in cell cycle exit during development (Goodyear and Richardson, 1997). At E14, the average mitochondrial branch length was shorter in proximal compared to distal HCs (Fig 2C). We attribute this to the higher overall network density and close proximity of multiple adjacent mitochondria in the high frequency region, which could not be separated at the imaging resolutions used. The further remodelling of mitochondrial dynamics in proximal compared to distal HCs between E10 and E14 indicates more extensive metabolic reprogramming in high frequency HCs. This is likely a consequence of the distinct physiological properties emerging between the two HC types at this time (Fettiplace and Nam, 2019).

**Figure 2:**
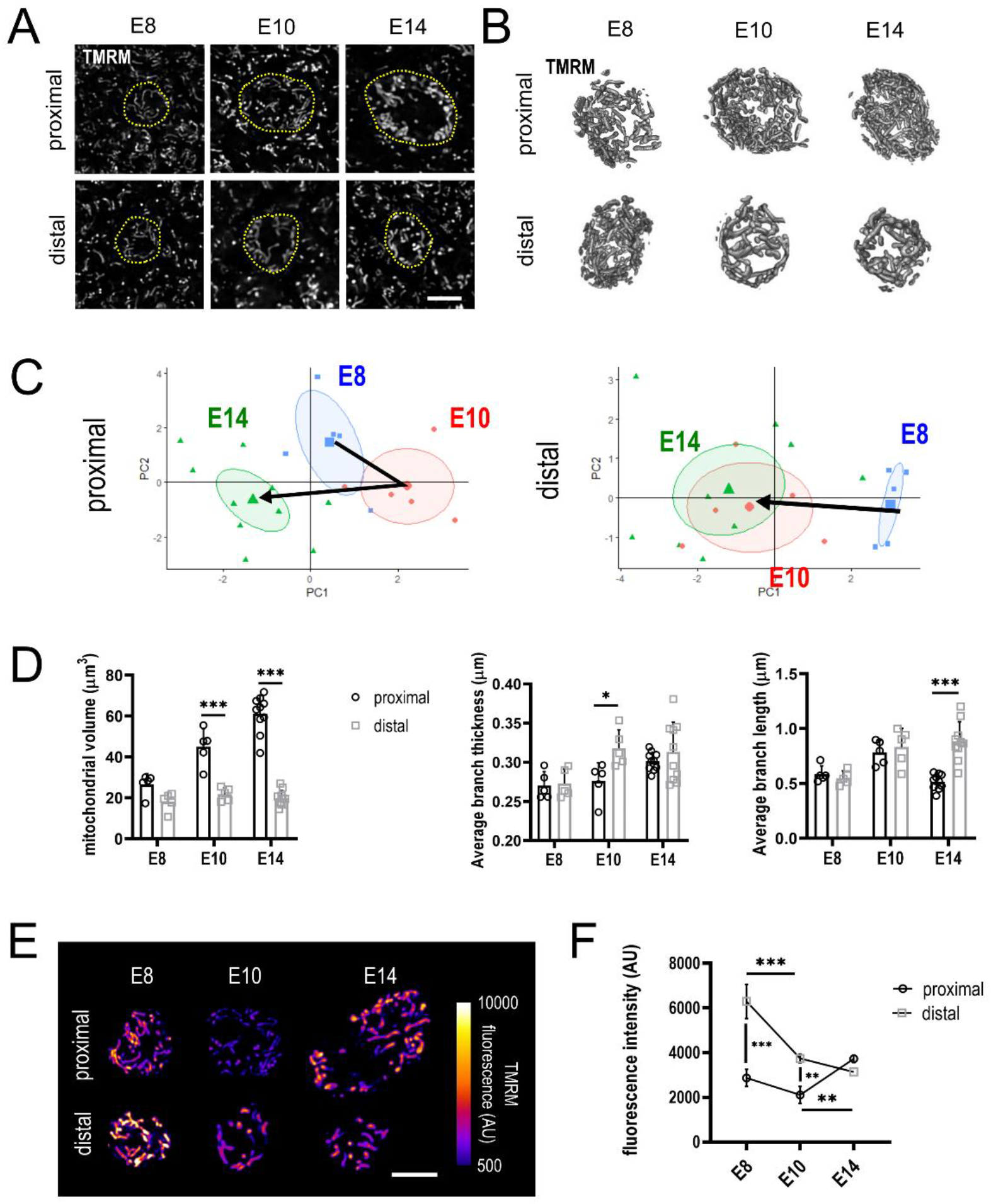
Mitochondrial morphology and membrane potential in developing HCs. **(A)** Example z-slices from explants at E8, E10 and E14 after loading with the mitochondrial dye TMRM. Z-stacks of the supranuclear mitochondrial population were acquired using TMRM fluorescence. HCs are delineated by yellow dashes. Scale = 5 µm. **(B)**3D models of supranuclear mitochondrial networks within HCs. Images were processed for noise reduction and edge enhancement. **(C)**Individual PCA plots of proximal and distal HC mitochondrial morphology, as observed from live (Airyscan) microscopy of TMRM-loaded BP explants dissected at E8, E10 and E14. Mitochondrial morphology in proximal HCs differs significantly between E8, E10 and E14 (circles – 95% confidence intervals), compared to distal HCs where mitochondrial morphology at E10 and E14 is comparable. Individual measurements which explain the variables contributing to PCs 1 and 2 are detailed in (D). **(D)**Quantification of individually measured mitochondrial shape descriptors. Data are mean ± SD, n = 6, 6, 11 HCs per timepoint. ^*^ p ≤ 0.05, ^***^ p ≤ 0.001, 2-way ANOVA followed by Sidak’s multiple comparisons. **(E)**Airyscan images of mitochondria from proximal and distal HCs in explants loaded with 350 nM TMRM at E8, E10 and E14. Images are single confocal slices of HC mitochondria demonstrating changes in HC mitochondrial membrane potential throughout development (E8 – E14). Scale bar = 5 µm. **(F)**At E8 and E10, mitochondria in distal HCs have a significantly higher membrane potential to those in proximal HCs. By E14, mitochondrial membrane potentials have equilibrated and become similar. Data displayed as mean ± SEM, n = 3 HCs from 2 independent biological replicates. ^**^ p ≤ 0.01, ^***^ p ≤ 0.001, 2-way ANOVA; Sidak’s multiple comparisons, F = 46.85, DF = 18.

To investigate how mitochondrial activity scales with mitochondrial content and HC cell size at different tonotopic positions, we quantified the fluorescence intensity of TMRM-loaded mitochondria, itself a read-out of the mitochondrial membrane potential (ΔΨm) and thus mitochondrial activity (Duchen et al., 2003). Unexpectedly, we observed that despite similar mitochondrial morphology at E8, distal HC mitochondria had a higher TMRM fluorescence and thus ΔΨm compared to proximal HCs (Fig 2E, F), which was maintained until E10 (Fig 2E, F). Although increased branch length is typically associated with increased mitochondrial activity, our observations suggest that in developing HCs, mitochondrial morphology and membrane potential are not tightly coupled. Furthermore, by E14, when proximal and distal mitochondria have largely different morphologies, ΔΨm had equalised between the two HC types (Fig 2B). These findings suggest at least in HCs, ΔΨm and thus mitochondrial activity is independent of their dynamics. The reduction in proximal ΔΨm at E8 and E10 could result from reduced activity in the mitochondrial TCA cycle. This may occur due to the increased glucose flux into the pentose phosphate pathway in the proximal region, which in turn reduces mitochondrial pyruvate uptake (O’Sullivan et al., 2023).

To investigate whether proximal-to-distal differences in the balance between fusion and fission regulators might underlie the gradient in mitochondrial morphology, we estimated expression levels of the fusion protein MFN1 and the active phosphorylated (Ser616) form of the fission GTPase DNM1L at E8, E10 and E14 using quantitative fluorescence immunohistochemistry. We detected no statistical differences in HC MFN1 immunoreactivity at any developmental stage (Fig S2A, B). In contrast, pDNM1L immunoreactivity was significantly higher in distal compared to proximal HCs at E8 (Fig S2C, D), and subsequently decreased in distal HCs between E8 and E14. The distal-to-proximal gradient in pDNM1L would be consistent with a bias towards fusion in the mitochondrial network of proximal HCs early in the growth window. The reduced distal pDNM1L immunoreactivity at E14 suggests a flattening of the distal-to-proximal fusion gradient towards the end of the growth window.

Previous work suggested that biasing cells towards fusion increases their size (Miettinen and Björklund, 2016). We directly tested this hypothesis using a chick fibroblast cell-line. Chicken DF-1 fibroblasts were transfected with plasmids containing a customisable CRISPR-Tol2 construct expressing a sgRNA guide and GFP reporter and Cas9 (Daudet et al., 2022) that allowed the expression of *MFN1* and *DNM1L* to be manipulated (Fig. 3A). *DNM1L* knockout (KO) fibroblasts exhibited higher mitochondrial volume (Fig 3B) and larger cell areas compared to control fibroblasts (Fig 3C), confirming that a bias towards fusion favours a larger cell size. When the morphological characteristics of mitochondria from *DNM1L* KO and control fibroblasts were plotted onto the PCA analysis conducted for E8 and E10 HCs, the difference in morphological trajectory between *DNM1L* KO and control fibroblasts matched those observed between proximal and distal HCs at E10 (Fig 3D). This supports the hypothesis that biasing mitochondrial dynamics towards fusion in proximal HCs could drive the tonotopic differences in supranuclear growth.

**Figure 3:**
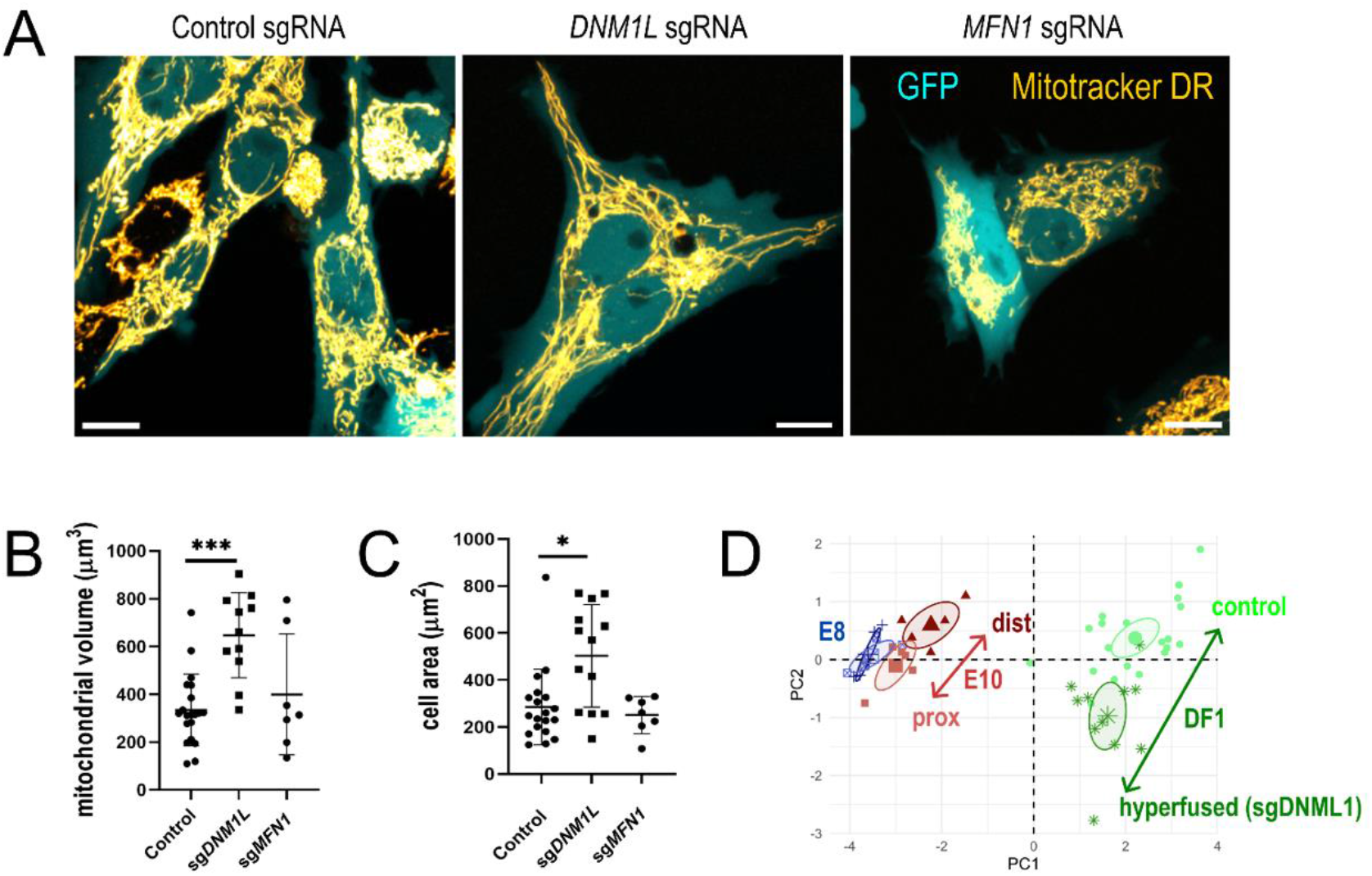
CRISPR-Cas9 knockout of fusion and fission regulators in DF-1 cells. **(A)** Maximum intensity projections of confocal z-stacks of GFP (cytosol) and MitoTracker DR (mitochondria) in DF-1 cells. Scale = 5 µm. **(B)** Total mitochondrial volume was increased in cells transfected with sgDNM1L compared to controls. No difference was observed in sgMFN1-transfected cells. Data are Mean ± SD, ^*^ p < 0.05, ^***^ p < 0.001 Kruskal Wallis, Sidak’s multiple comparisons; statistic = 14.03, DF = 26. **(C)** Cell area was increased in DF-1 cells transfected with sgDNM1L compared to controls. No difference was observed in sgMFN1 cells. Data are Mean ± SD, ^*^ p < 0.05, Kruskal Wallis, Sidak’s multiple comparisons; statistic = 8.742, DF = 26. **(D)** Morphometric analysis of control and sgDNM1L cell mitochondria reveals that their characteristics differ along the same axis as E10 proximal and distal HCs.

The cellular processes regulated by mitochondrial dynamics are determined by the rate of fission-fusion and how quickly this equilibrium can adjust (Spurlock et al., 2020). This rate differs between mitotic and postmitotic cells as well as across cell types. We therefore sought to functionally examine a role for mitochondrial dynamics in HC positional identity using pharmacological inhibitors to manipulate fusion and fission (Fig 4A) during the period of postmitotic cell growth between E9 and E12 (Fig 4B). Treatment with the fission inhibitor mdivi-1 (Cassidy-Stone et al., 2008) or the fusion promoter M1 (Wang et al., 2012) alone did not induce notable changes in the HC LSA (Fig 4C, D). However, co-treatment with mdivi-1 and M1 increased the distal HC LSA to levels comparable to those of proximal HCs (Fig 3B, C). Inhibition of fusion with MYLS22, an inhibitor of OPA1 (Baek et al., 2023; Herkenne et al., 2020), reduced HC size in the proximal BP region to levels comparable to distal HCs. Taken together, these data suggest that blocking fusion is sufficient to prevent supranuclear growth in proximal HCs and that a proximal-to-distal gradient in mitochondrial fusion may establish the tonotopic gradient in HC size. Although M1 has been identified as a fusion promoter, it does not act directly on any of the core fission-fusion machinery. Instead, M1 regulates activity of the mitochondrial ATP-synthase, whose fusion-promoting activity is blocked by oligomycin (Wang et al., 2012). The reduced effect of M1 on cell size in BP explants could be attributed to the limited response of HCs to oligomycin.

**Figure 4:**
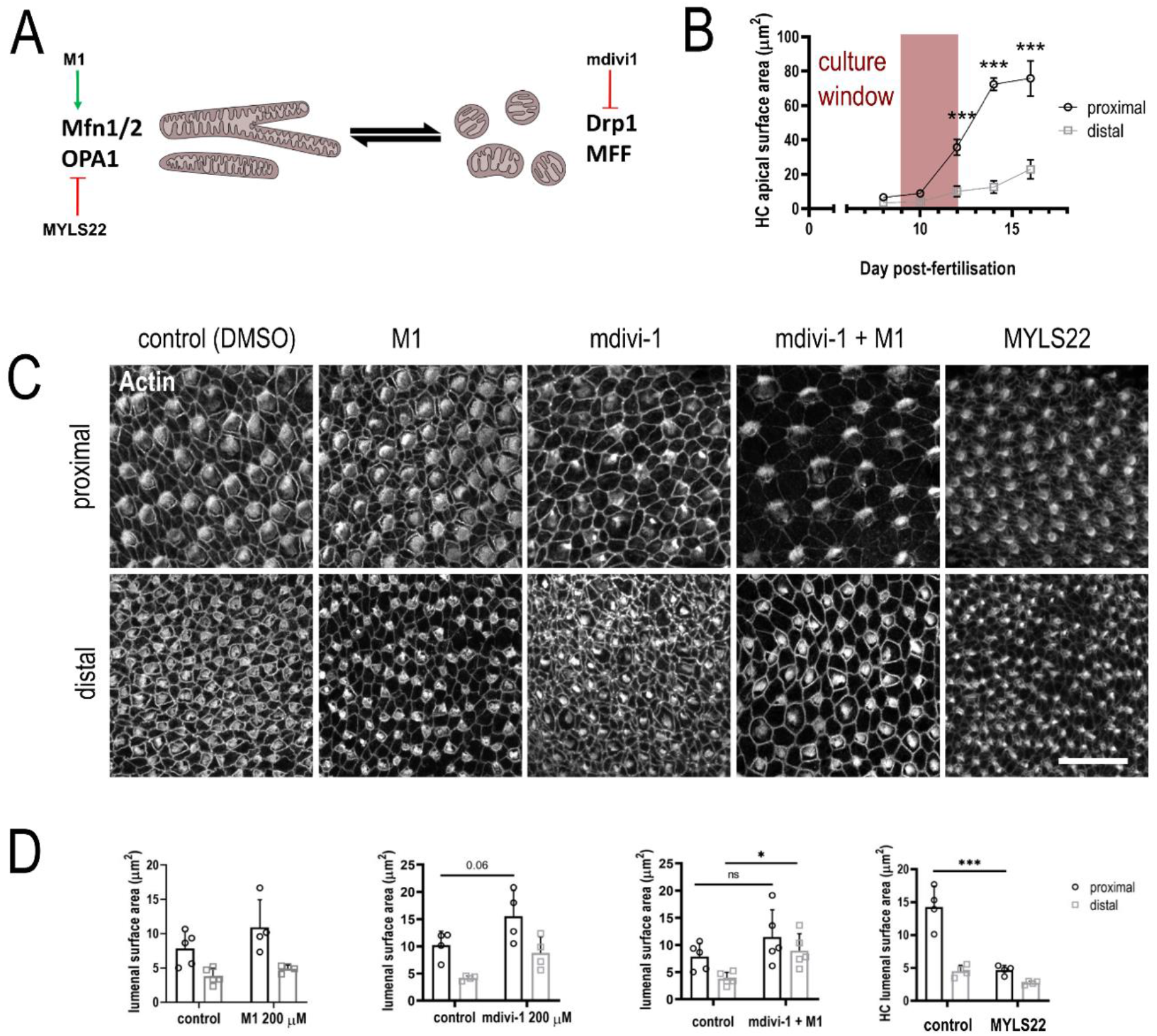
Functional interrogation of mitochondrial dynamics in the developing cochlea. Cochleae were dissected from embryonic chickens at E9. The sensory epithelium was isolated and cultured ex vivo for 4 days with small molecule modulators of mitochondrial dynamics. HC apical surface area was delineated using phalloidin staining. **(A)** Illustration depicting the site of action of different small molecule modulators. Mdivi-1 inhibits DNM1L and therefore fission. M1 promotes fusion. MYLS22 inhibits IMM fusion via OPA1. **(B)** Quantification of in vivo changes of HC size (E8-E16) reproduced from Figure 1 B. The time window for explant culture experiments is indicated by the pink band. **(C)** Effects of mitochondrial dynamics modulators on HC apical surface area. By themselves, M1 and mdivi-1 have no effects. However, M1 and mdivi1 in combination results in a large increase in distal HC surface area. Conversely, treatment with MYLS22 reduced apical HC surface area. Scale bar = 20 µm. **(D)** Quantification of modulator effects shown in **(C)**, n = 4, 4, 5, 4 independent biological replicates. Data are mean ± SEM. mdivi1+M1: 2-way ANOVA; Sidak’s multiple comparisons, t = 2.491, DF = 16, ^*^ p < 0.05. MYLS22: 2-way ANOVA; Sidak’s multiple comparisons, t = 8.01, DF = 12, p < 0.0001.

Overall, our findings suggest that differences mitochondrial fusion in the supranuclear region of HCs are a driver, rather than a consequence, of the larger high-frequency HC size. These morphological differences manifest as distinct surface-volume (SV) ratios at different tonotopic positions which determine the physiological properties of a HC (Fettiplace and Nam, 2019). Intracellular signalling pathways associated with cellular growth such as mTORC1 may be involved in regulating mitochondrial morphology in specific regions of the HC but are currently unexplored (Lloyd, 2013; Morita et al., 2017). The developmental regulation of asymmetric HC growth provides an informative model for investigating localised mTOR dependent activation of translation, mitochondrial dynamics and asymmetric growth. The identification of mitochondrial fusion as a regulator of HC size represents another key function of mitochondria in the ear, and demonstrates how insights from cell lines in previous literature (Miettinen and Björklund, 2016) and our own experiments can be translated to understanding cell size regulation in complex tissues.

## Materials and Methods

### Experimental animals

Fertilised chicken eggs (Dekalb white breed) were obtained from a commercial supplier (Medeggs LTD, Fakenham) and incubated at 37°C prior to experiments. All procedures were performed in accordance with the United Kingdom Animals (Scientific Procedures) Act (1986).

### CRISPR-Cas9 knockout constructs

pCAGG-NLS-Cas9-NLS (expressing nuclear localised *S. pyogenes* Cas9 under the expression of CAGC promoter) was a gift from Marianne Bronner (Addgene plasmid # 99138; http://n2t.net/addgene:99138; RRID: Addgene_99138). Tol2-U6.3-sgRNA-GFP was made by VectorBuilder expressing within a Tol2 transposon a cytoplasmic GFP reporter under the control of GAGC promoter and sgRNA guide expressed under chick specific U6.3 promoter. sgRNA guides sequences were designed using https://genome-euro.ucsc.edu/ with the March 2018 GRCg6a/galGal6 Gallus gallus genome release and chosen with following parameters: MIT specificity score > 80%, Efficiency Score >80% (Doench et al., 2016) and Out-of-Frame Score >65 (Bae et al., 2014). sgRNA were made by annealing pairs of synthesised complementary oligos (Eurofins-MWG) containing 20 base pair sgRNA guide sequences with additional 4 nucleotide overhangs (GGAT on forward, AAAC on reverse) for directional cloning into Tol2-U6.3-sgRNA-GFP at BsaI sites. sgRNA sequences are as follows: for *DNM1L* (GTGTTTTCCGACCATCCTCTG) targeting exon 2. For *MFN1* (GAGAAGAAGAGCGTCAAGGT) targeting exon 3. For non-targeting control (GCACTGCTACGATCTACACC).

### Transfection of DF-1 cells with CRISPR-Cas9 plasmid constructs

DF-1 fibroblasts were cultured in DMEM high glucose, GlutaMAX with sodium pyruvate (Thermo Fisher Scientific, 10569010) supplemented with 10% FBS at 5% CO^2^ 37 °C. DF-1 cells (50K per well) were plated into 24 µ-well imaging plates (ibidi -82426). Cells were transfected 24 hours later using lipofectamine 3000 (Thermo Fisher Scientific L3000008) with 1ug of plasmid NLS-Cas9-NLS and 0.5ug of appropriate Tol2-sgRNA -GFP plasmids, Cells were imaged live 72 hours after transfection. DF1 cells we purchased directly from ATTC (Chicken DF-1 fibroblast cells CRL-12203; ATCC) and screened routinely for any evidence of mycoplasma.

### Explant culture

The auditory sensory epithelium was dissected from chicken embryos at E9. Explants were adhered to glass bottomed Mattek dishes using 20% v/v CellTak solution according to the manufacturer’s instructions. Tissue explants were cultured for 96 hours in M199 medium supplemented with 2% bovine serum albumin and 10 mM HEPES at 37°C in 5% CO_2_. For small molecule treatments, stock solutions of MYLS22, mdivi-1 and M1 in DMSO were diluted in medium to their final concentrations. Medium for control treatments was supplemented with 0.2% DMSO (vehicle). Following the treatment protocol, explants were fixed for 20 minutes in 4% PFA in PBS pH 7.2.

### Histochemistry and Immunohistochemistry

Tissue was harvested from embryos for immunohistochemistry of MFN1 and phosphorylated (Ser616) DNM1L. The otic capsule was dissected from E8, E10 and E14 embryos and then fixed in 4% PFA in PBS pH 7.2 for one hour. The auditory sensory epithelium was dissected and permeabilised with 0.3% Triton X100. Blocking was performed with 10% goat serum in 0.3% Tween 20, before overnight incubation with unconjugated primary antibodies at 4°C (MFN1; Proteintech Polyclonal Rb 13798-1-AP, lot no. 00107635 1:100. pDNML1; Bioss Polyclonal Rb bs-12702R 1:100, lot no. BAO1271119. Calbindin: Abcam Monoclonal Ms ab82812-1001 1:75, lot no. 1027333-1). Tissue was stained with fluorophore-conjugated secondary antibodies at room temperature for one hour (Thermo Fisher Scientific A11001 + A11035), before post-staining with DAPI and phalloidin-633 (Thermo Fisher Scientific A22284) to detect DNA and F-actin respectively. Finally, samples were mounted on 8-well slides with Prolong-gold mountant.

### Transmission Electron Microscopy

Cochleae were dissected and fixed in 2.5% Glutaraldehyde in 0.1 M cacodylate buffer, pH 7.4, and were then bisected into proximal and distal portions before post-fixation in 1% buffered osmium tetroxide. Following post-fixation, samples were dehydrated to 70% ethanol before *en bloc* staining with 1% uranyl acetate in 70% ethanol. Samples were further dehydrated in ethanol before substitution with 100% propylene oxide. The samples were then infiltrated with increasing concentration of TAAB 812 hard resin, and mounted in the resin which was cured according to manufacturer instructions.

70 nm ultra-thin sections were taken using a Reichert Ultracut ultramicrotome. Ultra-thin sections were then stained with 1% aqueous uranyl acetate and 1% Lead citrate before imaging at 8000X magnification in a Jeol 1400 electron microscope with a 120 kV beam energy.

### Live imaging

Auditory sensory epithelia were dissected at E8, E10 and E14 and loaded with 350 nM TMRM (Thermo Fisher Scientific T668) in L-15 medium for 1 hour at room temperature. E14 tissue was dissected and loaded a 1000X diluted stock solution of CellTrace Calcein red-orange AM as indicated by the manufacturer (Thermo Fisher Scientific C34851). Dye-loaded explants were imaged in Mattek dishes on a Zeiss 980 confocal microscope, using the 561 nm laser and Airyscan 2 detector. Z-stacks were acquired, with an optical slice thickness of 18 nm. Airyscan processing was performed to software recommended settings. DF-Fibroblasts were imaged 72 hours after transfection and loaded with 100 nM Mitotracker Deep Red - MTDR (Thermo Fisher Scientific M22426) for 30 minutes 37°C in DMEM. Cells were imaged on Zeiss 880 confocal using a 488 nm laser for GFP and 633 nm laser and Airyscan 2 detector for MTDR.

### Cell volume

Cell volume was estimated using a stereology approach. For each HC, a cross-section was manually annotated every 1 µm interval of the z-stack. Using the slice-thickness of 0.18 μm, the volume of each HC was estimated.

### Mitochondrial network morphology

Raw data files were processed in the Fiji distribution of ImageJ (Schindelin et al., 2012) (Fig S3). To ensure compatibility with ImageJ’s skeleton module, images were first resampled to achieve isotropic pixel size using the CLIJ2 3D resample operation (Haase et al., 2020).

Subsequently, background fluorescence was eliminated using “subtract background” using a rolling ball size of 50 pixels. An unsharp masking filter was then applied, followed by the application of a non-local means filter. Next, a difference of gaussian filter was applied using minimum and maximum sigma of 2 and 4 pixels, respectively. Overall, we found this processing pipeline productive for minimising background and enhancing mitochondrial edges, leading to greater consistency between segmentation of different datasets.

Mitochondria were labelled using 3D WEKA segmentation. First, manual segmentation of data on a single confocal slice and used to train a classifier. The classifier was then used to segment mitochondria from the rest of the data analysed.

Binary stacks were generated from the 3D WEKA segmentation output. Next, individual HCs were cropped from the binary image stacks using ImageJ’s ROI tool, such that a single binary stack was generated for each HC in the dataset. Skeletons were generated for each HC image using the Skeleton extension. The BoneJ plugin (Doube et al., 2010) was used to analyse the binary stack from each HC, whilst the Analyse Skeleton plugin was used to analyse skeleton stacks to output branching information. From these two outputs measurements of local mitochondrial network characteristics were made, consisting of average branch thickness, standard deviation of branch thickness, maximum branch thickness, average branch length, maximum branch length, longest shortest path, Euclidean distance (a measure of branch tortuosity), and branch aspect ratio. These local mitochondrial characteristics were measured in HCs, and their developmental trajectories were compared between proximal and distal regions and visualised using PCA. Mitochondrial networks were plotted along principal components 1 and 2, providing graphical representation of these developmental trajectories.

### Statistical analysis

Principal component analysis (PCA) was carried out in R and visualised using the factoextra package.

For statistical analysis of live imaging, histochemical and IHC experiments, data were tested for normality and Q-Q plots were visually inspected to determine application of parametric and non-parametric tests. A mixture of 2-way ANOVA, t-tests and Kruskal-Wallis tests with appropriate post-hoc tests were carried out in Prism 8 (Graphpad Software, Boston MA).

## Acknowledgements

We thank Professor Jonathan Gale and Professor Andy Forge for their constructive comments and feedback on earlier versions of the manuscript.

---

**Figure S1:**
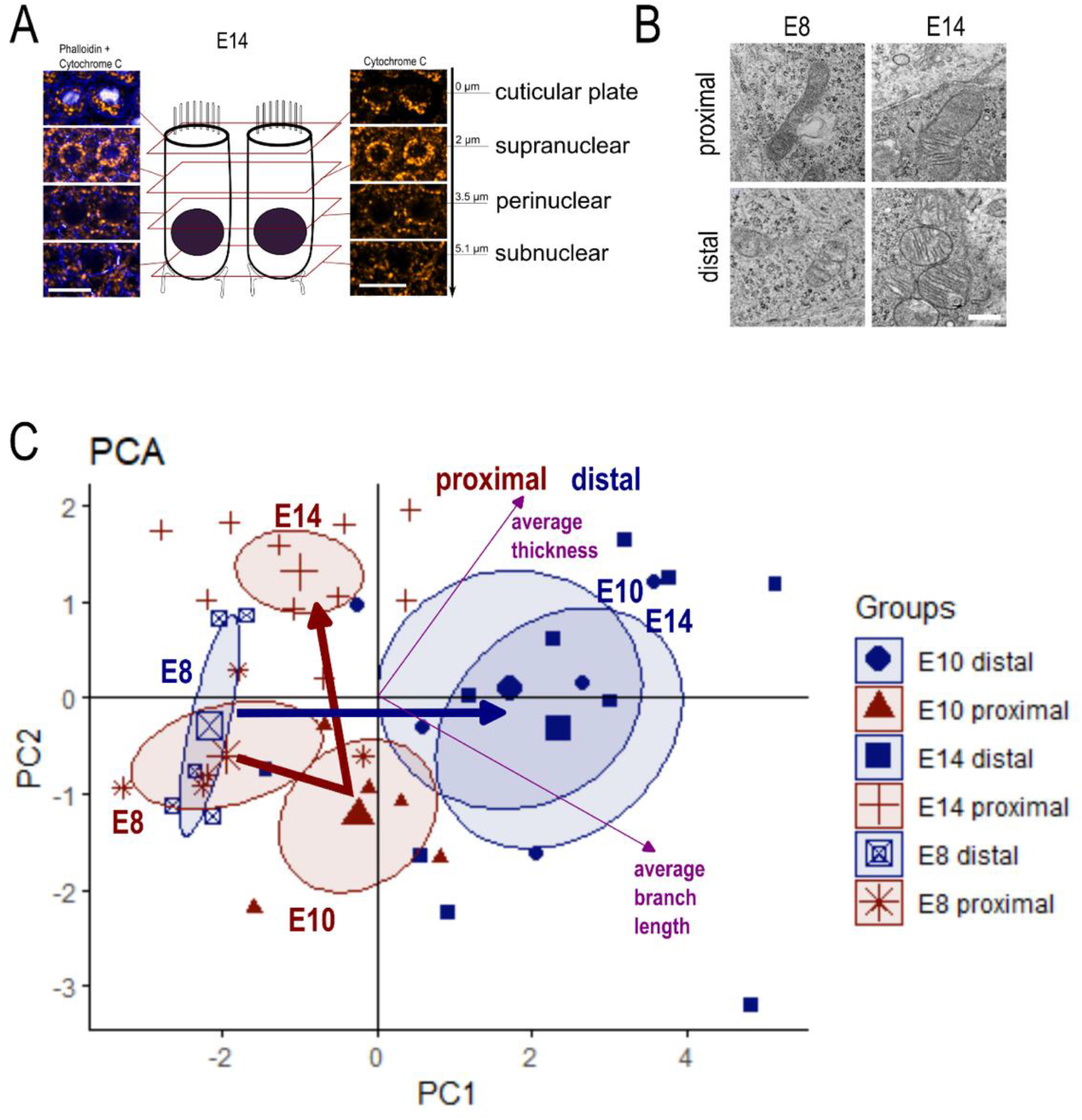
Mitochondrial morphology in cochlear HCs. **(A)** Immunohistochemistry of the mitochondrial antigen cytochrome c (orange), shown in combination with phalloidin (blue) at E14. Optical sections are taken from the different populations of mitochondria along the apico-basal axis of the HC. Scale = 10 µm. **(B)** TEM images of mitochondria within HCs at E8 and E14. Scale = 500 nm. **(C)** Principal component analysis (PCA) biplot of local mitochondrial network characteristics in HCs, including directionality of average thickness and branch length measurements (purple arrows). Each datapoint represents one HC sampled from distal (blue) or proximal (red) regions of the tissues. Larger points represent means, and ovals represent 95% confidence intervals. Arrows indicate the morphological trajectories of mitochondrial networks during HC development in proximal and distal regions.

**Figure S2:**
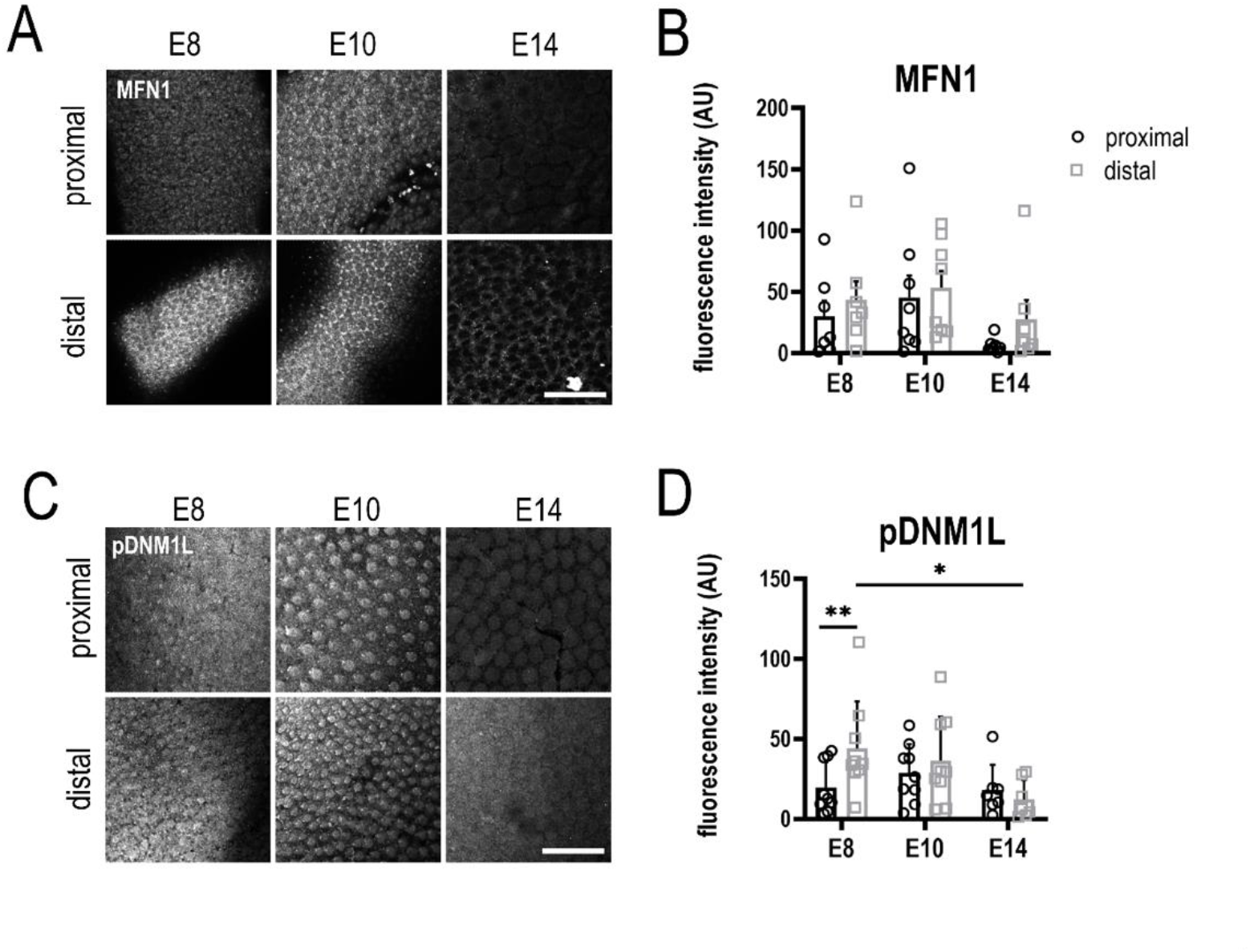
Expression of mitochondrial dynamics regulators in developing HCs. **(A)** Maximum intensity z-projection showing MFN1 immunoreactivity in cells of the BP sensory epithelium. Scale = 20 µm. **(B)** MFN1 immunoreactivity was consistent between proximal and distal regions at each timepoint. However, expression decreased in the proximal region between E10 and E14. Data are mean ± SEM, 15-40 HCs from 7 embryos, 2-way ANOVA. **(C)**Maximum z-projection showing pDNM1L immunoreactivity in the BP sensory epithelium. Scale = 20 µm. **(D)**pDNM1L immunoreactivity was significantly increased in distal HCs at E8, but then decreased between E8 and E14. Data are mean ± SEM, 15-40 HCs from 7 embryos, ^*^ p < 0.05, ^**^ p < 0.01, 2-way ANOVA: Sidak’s multiple comparisons, t = 3.385, DF = 22.

**Figure S3:**
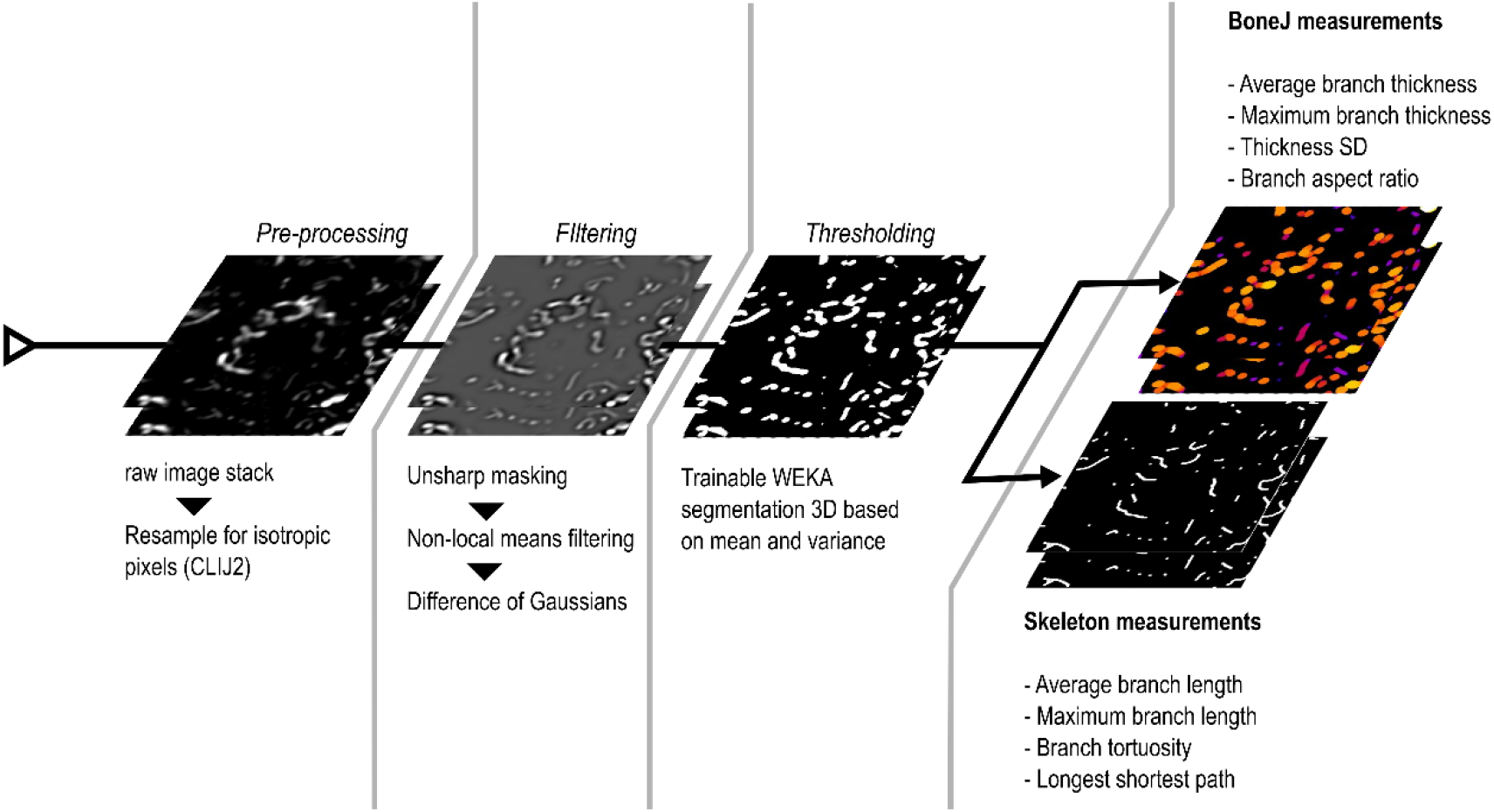
Image processing and analysis workflow for measurement of mitochondrial morphology.

## References

Afzal, N., Lederer, W. J., Jafri, M. S. and Mannella, C. A. (2021). Effect of crista morphology on mitochondrial ATP output: A computational study. Curr Res Physiol 4, 163–176.

Bae, S., Kweon, J., Kim, H. S. and Kim, J.-S. (2014). Microhomology-based choice of Cas9 nuclease target sites. Nat Methods 11, 705–706.

Baek, M. L., Lee, J., Pendleton, K. E., Berner, M. J., Goff, E. B., Tan, L., Martinez, S. A., Mahmud, I., Wang, T., Meyer, M. D., et al. (2023). Mitochondrial structure and function adaptation in residual triple negative breast cancer cells surviving chemotherapy treatment. Oncogene 42, 1117–1131.

Baum, T. and Gama, V. (2021). Dynamic properties of mitochondria during human corticogenesis. Development 148, dev194183.

Cassidy-Stone, A., Je, C., E, I., C, S., C, Y., T, K., Mj, K., Jt, S., Je, H., Dr, G., et al. (2008). Chemical inhibition of the mitochondrial division dynamin reveals its role in Bax/Bak-dependent mitochondrial outer membrane permeabilization. Developmental cell 14,.

Cervantes-Silva, M. P., Cox, S. L. and Curtis, A. M. (2021). Alterations in mitochondrial morphology as a key driver of immunity and host defence. EMBO Reports 22,.

Chang, C.-R., Manlandro, C. M., Arnoult, D., Stadler, J., Posey, A. E., Hill, R. B. and Blackstone, C. (2010). A Lethal de Novo Mutation in the Middle Domain of the Dynamin-related GTPase Drp1 Impairs Higher Order Assembly and Mitochondrial Division. J Biol Chem 285, 32494–32503.

Chen, H., Detmer, S. A., Ewald, A. J., Griffin, E. E., Fraser, S. E. and Chan, D. C. (2003). Mitofusins Mfn1 and Mfn2 coordinately regulate mitochondrial fusion and are essential for embryonic development. J Cell Biol 160, 189–200.

Cogliati, S., Frezza, C., Soriano, M. E., Varanita, T., Quintana-Cabrera, R., Corrado, M., Cipolat, S., Costa, V., Casarin, A., Gomes, L. C., et al. (2013). Mitochondrial cristae shape determines respiratory chain supercomplexes assembly and respiratory efficiency. Cell 155, 160–171.

Daudet, N., Żak, M., Stole, T. and Terry, S. (2022). Genetic Manipulation of the Embryonic Chicken Inner Ear. In Developmental, Physiological, and Functional Neurobiology of the Inner Ear (ed. Groves, A. K.), pp. 59–75. New York, NY: Springer US.

Doench, J. G., Fusi, N., Sullender, M., Hegde, M., Vaimberg, E. W., Donovan, K. F., Smith, I., Tothova, Z., Wilen, C., Orchard, R., et al. (2016). Optimized sgRNA design to maximize activity and minimize off-target effects of CRISPR-Cas9. Nat Biotechnol 34, 184–191.

Doube, M., Kłosowski, M. M., Arganda-Carreras, I., Cordelières, F. P., Dougherty, R. P., Jackson, J. S., Schmid, B., Hutchinson, J. R. and Shefelbine, S. J. (2010). BoneJ: Free and extensible bone image analysis in ImageJ. Bone 47, 1076–1079.

Duchen, M. R., Surin, A. and Jacobson, J. (2003). Imaging mitochondrial function in intact cells. Methods Enzymol 361, 353–389.

Fettiplace, R. and Nam, J.-H. (2019). Tonotopy in calcium homeostasis and vulnerability of cochlear hair cells. Hearing Research 376, 11–21.

Fu, W., Liu, Y. and Yin, H. (2019). Mitochondrial Dynamics: Biogenesis, Fission, Fusion, and Mitophagy in the Regulation of Stem Cell Behaviors. Stem Cells International 2019, 9757201.

Goodyear, R. and Richardson, G. (1997). Pattern Formation in the Basilar Papilla: Evidence for Cell Rearrangement. J. Neurosci. 17, 6289–6301.

Haase, R., Royer, L. A., Steinbach, P., Schmidt, D., Dibrov, A., Schmidt, U., Weigert, M., Maghelli, N., Tomancak, P., Jug, F., et al. (2020). CLIJ: GPU-accelerated image processing for everyone. Nat Methods 17, 5–6.

Herkenne, S., Ek, O., Zamberlan, M., Pellattiero, A., Chergova, M., Chivite, I., Novotná, E., Rigoni, G., Fonseca, T. B., Samardzic, D., et al. (2020). Developmental and Tumor Angiogenesis Requires the Mitochondria-Shaping Protein Opa1. Cell Metabolism 31, 987–1003.e8.

Iwata, R., Casimir, P. and Vanderhaeghen, P. (2020). Mitochondrial dynamics in postmitotic cells regulate neurogenesis. Science 369, 858–862.

Ji, W.-K., Chakrabarti, R., Fan, X., Schoenfeld, L., Strack, S. and Higgs, H. N. (2017). Receptor-mediated Drp1 oligomerization on endoplasmic reticulum. J Cell Biol 216, 4123–4139.

Khacho, M., Clark, A., Svoboda, D. S., Azzi, J., MacLaurin, J. G., Meghaizel, C., Sesaki, H., Lagace, D. C., Germain, M., Harper, M.-E., et al. (2016). Mitochondrial dynamics impacts stem cell identity and fate decisions by regulating a nuclear transcriptional program. Cell Stem Cell 19, 232–247.

Kitami, T., Logan, D. J., Negri, J., Hasaka, T., Tolliday, N. J., Carpenter, A. E., Spiegelman, B. M. and Mootha, V. K. (2012). A Chemical Screen Probing the Relationship between Mitochondrial Content and Cell Size. PLoS One 7, e33755.

Lippe, W. R. (1995). Relationship between frequency of spontaneous bursting and tonotopic position in the developing avian auditory system. Brain Res 703, 205–213.

Lloyd, A. C. (2013). The Regulation of Cell Size. Cell 154, 1194–1205.

Mann, Z. F. and Kelley, M. W. (2011). Development of tonotopy in the auditory periphery. Hear Res 276, 2–15.

Mann, Z. F., Thiede, B. R., Chang, W., Shin, J.-B., May-Simera, H. L., Lovett, M., Corwin, J. T. and Kelley, M. W. (2014). A gradient of Bmp7 specifies the tonotopic axis in the developing inner ear. Nat Commun 5, 3839.

Miettinen, T. P. and Björklund, M. (2016). Cellular Allometry of Mitochondrial Functionality Establishes the Optimal Cell Size. Dev Cell 39, 370–382.

Morita, M., Prudent, J., Basu, K., Goyon, V., Katsumura, S., Hulea, L., Pearl, D., Siddiqui, N., Strack, S., McGuirk, S., et al. (2017). mTOR Controls Mitochondrial Dynamics and Cell Survival via MTFP1. Molecular Cell 67, 922–935.e5.

O’Sullivan, J. D., Blacker, T. S., Scott, C., Chang, W., Ahmed, M., Yianni, V. and Mann, Z. F. (2023). Gradients of glucose metabolism regulate morphogen signalling required for specifying tonotopic organisation in the chicken cochlea. eLife 12, e86233.

Patten, D. A., Wong, J., Khacho, M., Soubannier, V., Mailloux, R. J., Pilon-Larose, K., MacLaurin, J. G., Park, D. S., McBride, H. M., Trinkle-Mulcahy, L., et al. (2014). OPA1-dependent cristae modulation is essential for cellular adaptation to metabolic demand. EMBO J 33, 2676–2691.

Posakony, J., England, J. and Attardi, G. (1977). Mitochondrial growth and division during the cell cycle in HeLa cells. J Cell Biol 74, 468–491.

Schindelin, J., Arganda-Carreras, I., Frise, E., Kaynig, V., Longair, M., Pietzsch, T., Preibisch, S., Rueden, C., Saalfeld, S., Schmid, B., et al. (2012). Fiji: an open-source platform for biological-image analysis. Nat. Methods 9, 676–682.

Si, F., Brodie, H., Gillespie, P. G., Vazquez, A. E. and Yamoah, E. N. (2003). Developmental Assembly of Transduction Apparatus in Chick Basilar Papilla. J. Neurosci. 23, 10815–10826.

Son, E. J., Ma, J.-H., Ankamreddy, H., Shin, J.-O., Choi, J. Y., Wu, D. K. and Bok, J. (2015). Conserved role of Sonic Hedgehog in tonotopic organization of the avian basilar papilla and mammalian cochlea. PNAS 112, 3746–3751.

Song, M., Mihara, K., Chen, Y., Scorrano, L. and Gerald W Dorn, I. I. (2015). Mitochondrial fission and fusion factors reciprocally orchestrate mitophagic culling in mouse hearts and cultured fibroblasts. Cell metabolism 21, 273.

Spurlock, B., Tullet, J., Hartman, J. L. and Mitra, K. (2020). Interplay of mitochondrial fission-fusion with cell cycle regulation: Possible impacts on stem cell and organismal aging. Exp Gerontol 135, 110919.

Thiede, B. R., Mann, Z. F., Chang, W., Ku, Y.-C., Son, Y. K., Lovett, M., Kelley, M. W. and Corwin, J. T. (2014). Retinoic acid signalling regulates the development of tonotopically patterned hair cells in the chicken cochlea. Nat Commun 5, 3840.

Tilney, L. G. and Saunders, J. C. (1983). Actin filaments, stereocilia, and hair cells of the bird cochlea. I. Length, number, width, and distribution of stereocilia of each hair cell are related to the position of the hair cell on the cochlea. Journal of Cell Biology 96, 807–821.

Van Der Bliek, A. M., Shen, Q. and Kawajiri, S. (2013). Mechanisms of Mitochondrial Fission and Fusion. Cold Spring Harbor Perspectives in Biology 5, a011072–a011072.

Wang, D. J W., Gm, B., S, M., Rg, B., C, T. P Y. and Pg, S. (2012). A small molecule promotes mitochondrial fusion in mammalian cells. Angewandte Chemie (International ed. in English) 51.

Wiemerslage, L. and Lee, D. (2016). Quantification of Mitochondrial Morphology in Neurites of Dopaminergic Neurons using Multiple Parameters. J Neurosci Methods 262, 56–65.

Wong, E. D., Wagner, J. A., Gorsich, S. W., McCaffery, J. M., Shaw, J. M. and Nunnari, J. (2000). The dynamin-related GTPase, Mgm1p, is an intermembrane space protein required for maintenance of fusion competent mitochondria. J Cell Biol 151, 341–352.

Zehnder, T., Petrelli, F., Romanos, J., De Oliveira Figueiredo, E. C., Lewis, T. L., Déglon, N., Polleux, F., Santello, M. and Bezzi, P. (2021). Mitochondrial biogenesis in developing astrocytes regulates astrocyte maturation and synapse formation. Cell Rep 35, 108952.

